# SWIP mediates retromer-independent membrane recruitment of the WASH complex

**DOI:** 10.1101/2022.08.16.504073

**Authors:** V. Dostál, T. Humhalová, P. Beránková, O. Pácalt, L. Libusová

## Abstract

The pentameric WASH complex facilitates endosomal protein sorting by activating Arp2/3, which in turn leads to the formation of F-actin patches specifically on the endosomal surface. It is generally accepted that WASH complex attaches to the endosomal membrane via the interaction of its subunit FAM21 with the retromer subunit VPS35. However, we observe the WASH complex and F-actin present on endosomes even in the absence of VPS35. We show that the WASH complex binds to the endosomal surface in both a retromer-dependent and a retromer-independent manner. The retromer-independent membrane anchor is directly mediated by the subunit SWIP. Furthermore, SWIP can interact with a number of phosphoinositide species. Of those, our data suggest that the interaction with phosphatidylinositol-3,5-bisphosphate (PI(3,5)P_2_) is crucial to the endosomal binding of SWIP. Overall, this study reveals a new role of the WASH complex subunit SWIP and highlights the WASH complex as an independent, self-sufficient trafficking regulator.

**Summary:** Dostál et al. contradict the prevailing concept that WASH complex is principally recruited to the endosome via its interaction with the retromer. They show that the WASH complex binds to the endosomal membrane via its subunit SWIP, and this interaction can be prevented by removing phosphatidylinositol-3,5-bisphosphate from cells.

## Introduction

Endosomes are essential sorting hubs of the cell, receiving material from the plasma membrane and various intracellular reservoirs and sending each cargo molecule to its proper destination. This process – endosomal protein sorting – is vital for the maintenance of cellular homeostasis. It is regulated by an intricate network of proteins which mediate cargo binding, sorting to endosomal microdomains, membrane tubulation, and finally scission of a transport vesicle. An important role in all of these steps is played by F-actin which often forms dense patches adjacent to the endosomal membrane (Durrbach et al., 1996; Nakagawa and Miyamoto, 1998; Puthenveedu et al., 2010; Simonetti and Cullen, 2018).

Endosomal F-actin assembly is driven by the pentameric WASH complex (Derivery et al., 2009; Gomez et al., 2012). The complex is named after its Wiskott-Aldrich syndrome protein and SCAR homologue 1 (WASH1) subunit, which belongs to the WASP family of nucleation-promoting factors (NPF). WASH1 activates Arp2/3 through its VCA domain and hence promotes nucleation of F-actin assembly (Linardopoulou et al., 2007). Besides WASH1, the WASH complex consists of proteins FAM21, strumpellin, SWIP, and CCDC53 (Derivery et al., 2009). However, specific functions of these subunits are not well understood. The whole complex localizes to distinct membrane domains of early and sorting endosomes where it facilitates recycling of cargo towards the Golgi apparatus (Gomez and Billadeau, 2009) and to the plasma membrane (Zech et al., 2011).

Various membrane proteins depend on the WASH complex for their proper subcellular localization. These include glucose transporter 1 (GLUT1), cation-independent mannose 6-phosphate receptor (CI-MPR) (Gomez and Billadeau, 2009; Piotrowski et al., 2013), major histocompatibility complex II (Graham et al., 2014), integrin α5β1 (Zech et al., 2011) and a growing list of other important integral membrane proteins. WASH1 knockout is embryonically lethal in mice (Gomez et al., 2012), and point mutations in WASH complex components are linked to serious human disorders including hereditary spastic paraplegia (Clemen et al., 2010) and autosomal recessive intellectual disability (Ropers et al., 2011). Clearly, ensuring the proper function of the WASH complex is essential for normal cell physiology.

To carry its function, the complex must localize to the outer surface of the endosomal membrane. The WASH complex associates with the retromer, a master regulator of endosomal recycling, and early studies postulated that the retromer is required for the membrane recruitment of the WASH complex to the endosomal membrane (Harbour et al., 2012; Jia et al., 2012). The interaction is mediated by the FAM21 subunit of the WASH complex and the VPS35 subunit of the retromer (Helfer et al., 2013). The VPS35 D620N mutation, which received considerable attention as it can manifest itself as a hereditary form of late-onset Parkinson disease (Zimprich et al., 2011), impairs the interaction with FAM21. It was suggested that this leads to WASH complex delocalization (Zavodszky et al., 2014). However, the ability of this mutation to displace the WASH complex from the endosomes has been disputed (McGough et al., 2014). In contrast to the theory of retromer-mediated recruitment of the WASH complex, FAM21 was found to associate with endosomes in a VPS35 knockout HeLa cell line (McNally et al., 2017), suggesting that there are additional factors attaching the WASH complex to the endosomal surface.

This additional membrane attachment of the WASH complex may be attributed to an interaction with an unknown membrane protein. However, despite several efforts to characterize the interactome of the WASH complex (Derivery et al., 2009; Freeman et al., 2014, p. 8; Ryder et al., 2013; Visweshwaran et al., 2018), no definite candidate for a membrane-anchoring protein has been described so far. Hepatocyte growth factor-regulated tyrosine kinase substrate (HRS) facilitates recruitment of WASH1 to the membrane but has not been shown to physically interact with it and localizes to a different endosomal microdomain (MacDonald et al., 2018). Recently, the discovery of the CCC and retriever complexes has furthered our understanding of the interaction network of the WASH complex (Phillips-Krawczak et al., 2015, p. 1). However, despite the homology of the retriever to the retromer complex, the retriever is in fact dependent on the WASH complex for its membrane localization and not vice versa (McNally et al., 2017). Therefore, there is currently no accepted surrogate for a WASH complex membrane-recruiting protein besides the retromer.

Alternatively, the WASH complex may harbor intrinsic phospholipid-binding ability. Indeed, a purified WASH complex is able to directly interact with artificial liposomes (Derivery et al., 2009). Moreover, the C-terminal end of the FAM21 subunit potently interacts with various phosphoinositide (PI) species and phosphatidylserine (PS) on a PIP strip overlay assay (Jia et al., 2010). Nevertheless, the binding to phospholipids could be mediated by any of the subunits of the WASH complex except for CCDC53, whose localization is entirely dependent on WASH1 (Gomez et al., 2012).

In this study, we show that the membrane binding of the WASH complex and its actin-polymerizing activity do not require the retromer-FAM21 interaction. Using a set of knockout cell lines for individual components of the WASH complex, we clearly determine that the SWIP subunit associates with endosomal membrane independently of FAM21 and helps recruit the rest of the WASH complex to the endosomal membrane.

## Results

### WASH complex membrane attachment and function do not require retromer

As the studies that examined the retromer-mediated WASH complex recruitment provided contradictory results, we decided to reinvestigate WASH complex localization in cells where the WASH-retromer link (Fig. 1A) is perturbed. To do so, we depleted one member of the retromer, protein VPS35, from the mammalian cell line U-2 OS using CRISPR/Cas9 (Fig. S1A).

**Figure 1.**
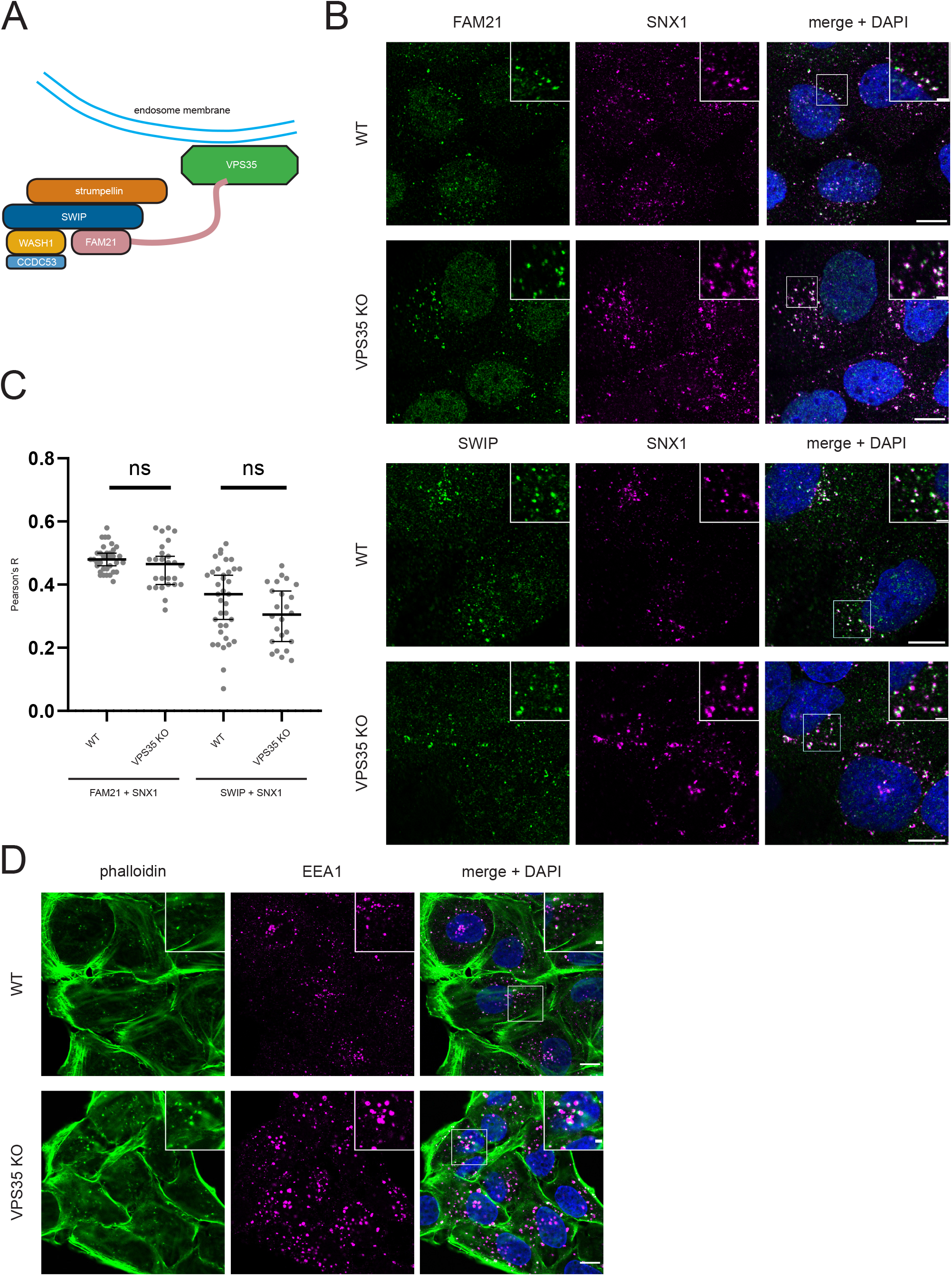
VPS35 in not required for the WASH complex endosomal localization and function. **(A)** Original model of retromer-mediated WASH complex attachment to endosomal membrane (adapted from Seaman et al., 2013). **(B)** Wild-type U-2 OS cells and VPS35 knockout U-2 OS cells were fixed and stained with antibodies targeting SNX1 and WASH complex subunits FAM21 or SWIP. Insets show WASH complex subunits localized to SNX1-positive vesicles. **(C)** Quantification of colocalization between SNX1 and FAM21 or SNX1 and SWIP in WT and VPS35 KO cells. Each dot represents one analyzed cell. Data from 3 independent experiments, >20 cells in total analyzed in each column (nuclear regions removed from analyzed ROIs). **(D)** WT and VPS35 KO cells were labeled with phalloidin and an anti-EEA1 antibody. The staining shows patches of endosomal actin in both WT and VPS35 KO cell lines. Scale bars: 10 μm (main images), 2 μm (insets). Quantitative data in graphs tested for normality with Shapiro-Wilk test, significance calculated using the unpaired t-test. Lines in graphs indicate median ± 95% confidence interval.

We then analyzed the WASH complex localization in this VPS35 knockout (KO) cell line. We immunostained FAM21, SWIP and WASH1 subunits in WT and VPS35 KO cell lines together with SNX1, a marker of the corresponding endosomal microdomain. SNX1-labeled endosomes are enlarged in the VPS35 KO cell line. However, the localization of the WASH complex subunits to SNX1-positive vesicles is unperturbed (Fig 1B–C and Fig. S1B). This confirms that the retromer is not essential for the membrane attachment of the WASH complex.

Therefore, we hypothesized that the perturbation of the WASH-retromer interaction influences the WASH complex function rather than its localization. Thus, we evaluated the actin-nucleating activity of the WASH complex in the absence of VPS35. We stained F-actin and early endosomes in WT and VPS35 KO using fluorescently labeled phalloidin and an anti-EEA1 antibody (Fig. 1D). Even though the depletion of VPS35 increased the size of the EEA1-labeled early endosomes, in line with SNX1 pattern, the endosomal actin patches were intact in the VPS35 KO cell line. This further confirms that the WASH complex does not depend solely on VPS35 to interact with the endosomal surface and carry out its function there.

### Function of the WASH complex is abolished by knockout of WASH1, FAM21, SWIP or strumpellin

We next sought to identify the specific subunits required for WASH complex localization and function. Since the CCDC53 subunit delocalizes from endosomes upon WASH1 KO in mammalian cells while the rest of the complex remains attached (Gomez et al., 2012), we excluded CCDC53 from our search. Using the CRISPR/Cas9 system in U-2 OS cells, we generated FAM21 KO, SWIP KO, strumpellin KO, and WASH1 KO cell lines. Validity of each cell line was confirmed by Western blotting (Fig. 2A) and sequencing of the affected regions.

**Figure 2.**
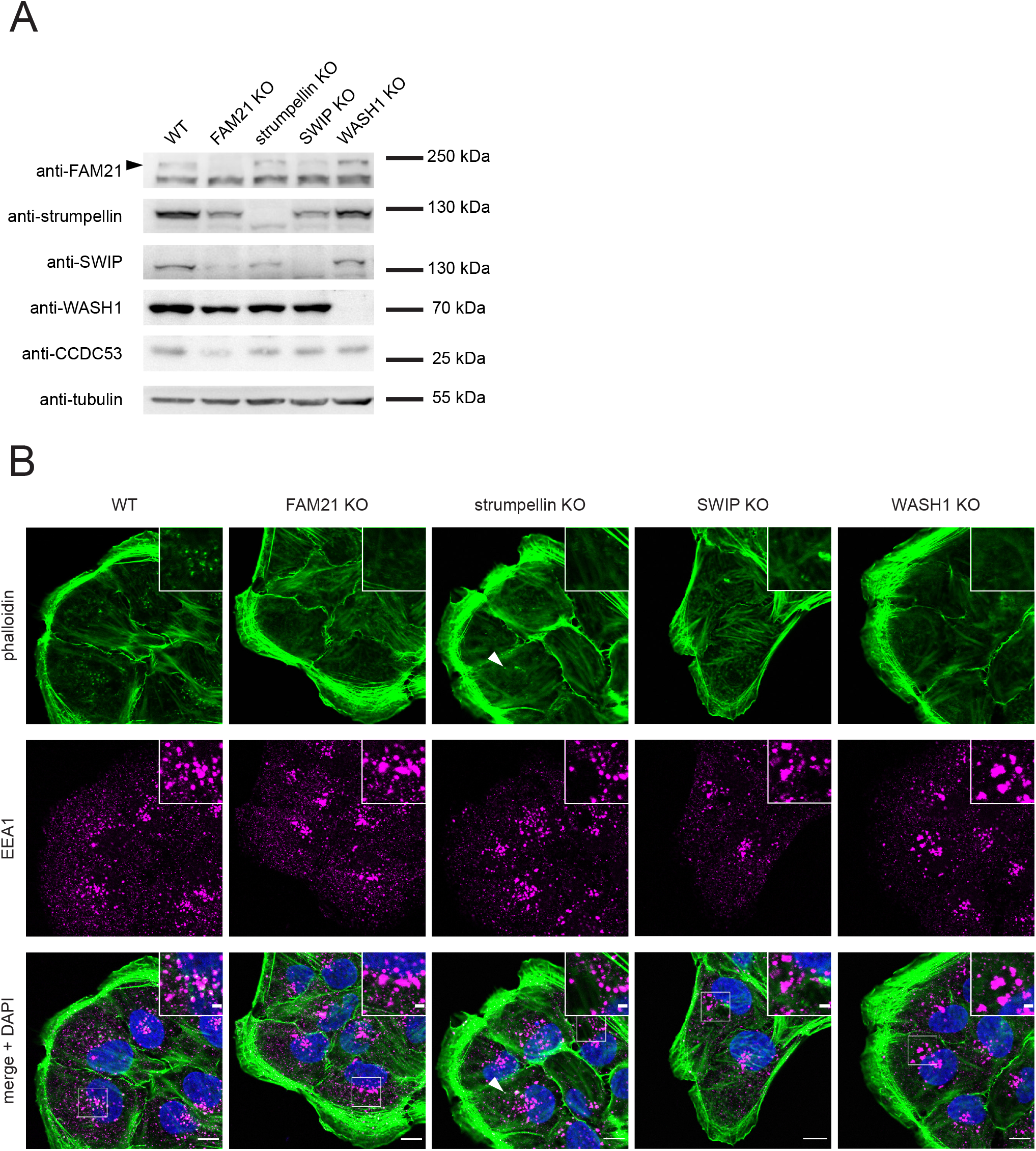
Perturbation of the WASH complex leads to loss of endosomal actin. **(A)** Endogenous protein levels in WASH complex knockout cell lines were analyzed by Western blotting followed by staining with antibodies targeting indicated proteins. Black arrow denotes a FAM21-specific band. **(B)** Indicated knockout cell lines were fixed and labeled with phalloidin-A488 and antibody targeting early endosomal protein EEA1. Insets are given for a detailed view of endosomal patches. White arrow in strumpellin KO denotes a possible remnant of endosomal patches in this cell line. Scale bars: 10 μm (main images), 2 μm (insets).

We then explored the functional implications of individual WASH complex subunit knockouts. First, expression of all WASH complex subunits was evaluated in each cell line. The cells were not only devoid of the targeted subunits, but also often displayed decreased protein levels of the remaining subunits of the complex (Fig. 2A), which is in line with previous studies (Gomez et al., 2012).

Intriguingly, the active subunit of the complex, WASH1 protein, was less susceptible to this effect, and its levels remained largely steady in all knockout cell lines. We thus wondered if WASH1 also retains its ability to promote F-actin nucleation in the absence of the other subunits. Co-staining by phalloidin and an anti-EEA1 antibody in these cell lines revealed that endosomal actin patches are absent upon knockout of WASH1, SWIP or FAM21 and largely diminished in the strumpellin KO cell line (Fig. 2B). Thus, even though the levels of WASH1 were virtually unchanged upon depletion of other WASH complex subunits, the protein lost its ability to drive actin polymerization on endosomes. This demonstrates the essentiality of each of the subunits, with the possible exception of strumpellin, where weak endosomal F-actin signal remains in some strumpellin KO cells. To confirm the specificity of the observed phenotypes, we sought to rescue expression of SWIP in the SWIP KO line. Reintroduction of EGFP-SWIP into SWIP KO cells recovered endosomal F-actin patches which co-localized with EGFP-SWIP signal (Fig. S2A).

We also analyzed how the absence of endosomal actin nucleation in WASH KO cells translated into the trafficking of GLUT1 transporter, a canonical cargo of the WASH complex. Unsurprisingly, GLUT1 is missorted from the plasma membrane in WASH1 KO, strumpellin KO, SWIP KO, and FAM21 KO cell lines and instead accumulates in endosomes (Fig. S2B).

### Subunits FAM21 and SWIP both link the WASH complex to endosomes

We then decided to identify which individual WASH complex subunits are essential for its endosomal recruitment. To do so, we analyzed localization of the remaining subunits in all generated WASH complex KO cell lines. Even though our previous experiments showed that the levels of FAM21, strumpellin, SWIP, and CCDC53 were diminished in the KO cell lines, the residual levels were sufficient for microscopic analysis. We immunostained the individual WASH complex subunits and VPS35 in each WASH complex KO cell line. VPS35, which almost perfectly co-localizes with WASH complex subunits in WT cells, served as a marker of the appropriate WASH-decorated microdomain of endosomes (Fig. 3 and S3).

**Figure 3.**
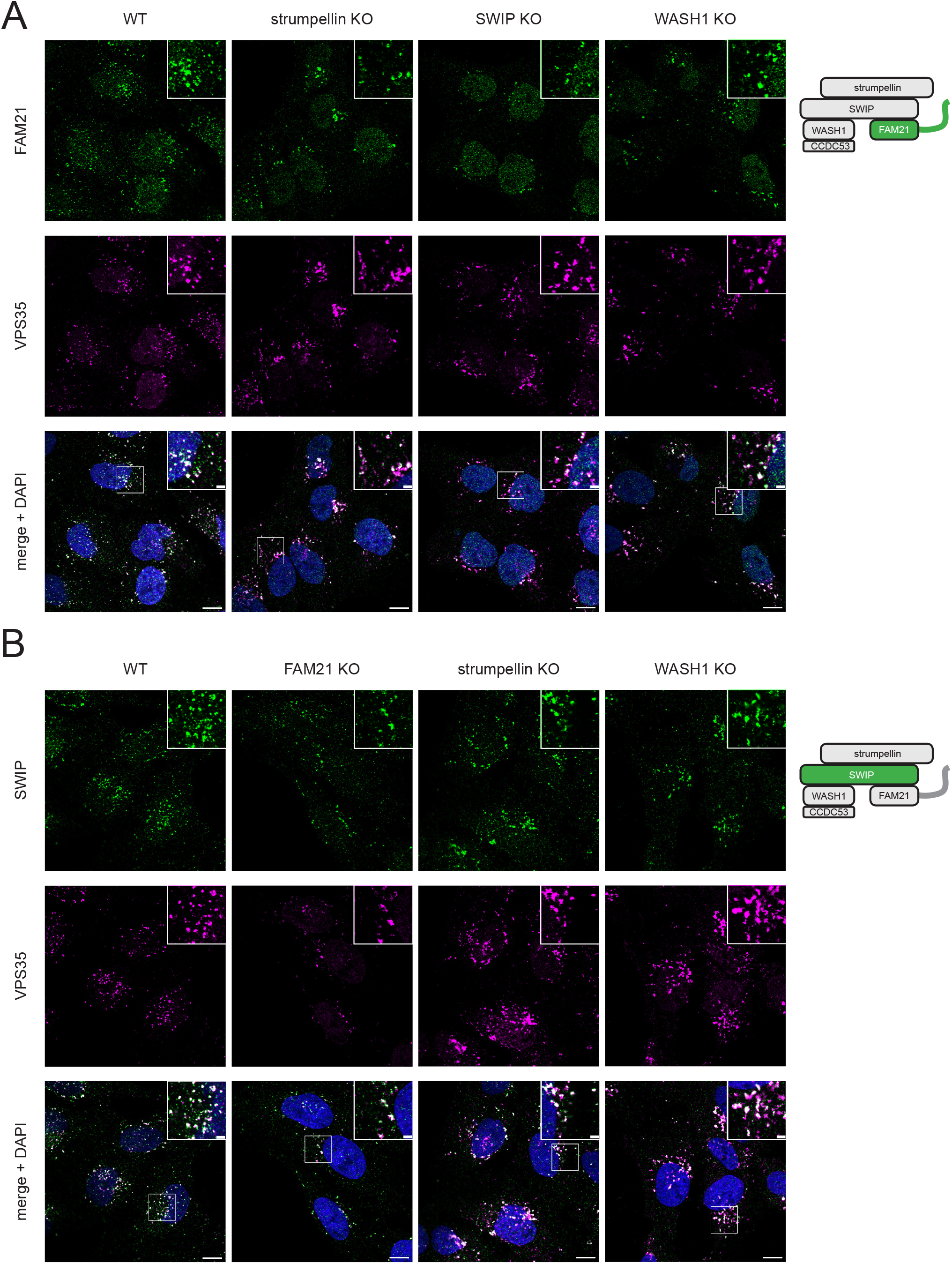
FAM21 and SWIP do not require other WASH complex subunits for their endosomal localization. Wild-type U-2 OS cells and indicated knockout cell lines were fixed and labeled with antibodies targeting VPS35 and FAM21 **(A)** or SWIP **(B)**. The staining shows that FAM21 and SWIP are attached to endosomes in all presented KO cell lines. Scale bars: 10 μm (main images), 2 μm (insets). Schematic model of the WASH complex with the labeled subunits in green illustrates the performed experiments.

Results of the microscopic analysis are summarized in Table 1. As a rule, FAM21 and SWIP never delocalize from the endosomes in response to any of the subunit knockouts (Fig. 3A–B). This result was expected for FAM21 due to its retromer-binding properties, but it is a novel observation for SWIP, which does not detach from membranes even in FAM21 KO cells. The remaining subunits are dependent on at least one other subunit for their localization. WASH1 and CCDC53 completely delocalize from the endosomes in FAM21 KO cells (Fig. S3A and S3C), while strumpellin only delocalizes in the SWIP KO cell line (Fig. S3B). CCDC53 delocalizes in WASH1 KO cells, confirming previous reports that CCDC53 binds to the complex via WASH1, and its localization is also partially hampered in SWIP KO cells.

**Table 1.** Localization analysis of individual WASH complex subunits. Table summarizing the results of localization analysis of individual WASH complex subunits in wild-type U-2 OS and cell lines lacking FAM21, Strumpellin, SWIP or WASH1 (see Fig. 3 and Fig. S3).

|  | WT | FAM21<br>KO | strum-<br>pellin<br>KO | SWIP<br>KO | WASH1<br>KO |
| --- | --- | --- | --- | --- | --- |
| FAM21 | vesicles | - | vesicles | vesicles | vesicles |
| strum-<br>pellin | vesicles | vesicles | - | not<br>localized | vesicles |
| SWIP | vesicles | vesicles | vesicles | - | vesicles |
| WASH1 | vesicles | not<br>localized | vesicles | vesicles | - |
| CCDC53 | vesicles | not<br>localized | vesicles | few<br>vesicles | not<br>localized |

Together, these results indicate that SWIP and FAM21 subunits are both independently able to interact with the endosomal surface. In contrast, the endosomal localization of strumpellin is dependent on SWIP while the localization of WASH1 and CCDC53 is most sensitive to the absence of FAM21, suggesting that the membrane attachment of these subunits follows either of the two main anchors – FAM21 or SWIP. However, partial dissociation of CCDC53 in SWIP KO indicates that SWIP is also involved in CCDC53 localization.

### FAM21 binds to the endosomes via SWIP and VPS35

The interaction between FAM21 and VPS35 represents the established link of the WASH complex to the endosomal membrane. In light of our data, we asked whether VPS35 and SWIP could work together to link FAM21 to the endosomal membrane. To investigate this, we decided to create a double KO cell line lacking VPS35 and SWIP, but all resulting cells died after several rounds of cell division. As an alternative solution, we analyzed the localization of FAM21 in SWIP KO cells which were transiently depleted of VPS35 by siRNA transfection. Under these conditions, FAM21 was not detectable on endosomes (Fig. S4A). This could be due to extremely low protein levels of FAM21 (Fig. S4B), likely resulting from the absence of its two important interaction partners. We therefore aimed to determine whether the loss of the FAM21 endosomal signal is due to delocalization or a decreased amount of the protein. To address this issue, we increased cellular FAM21 levels by expressing FAM21-EGFP in the SWIP KO siVPS35 cells and counter-stained them with antibody targeting SNX1 (Fig. 4A–C). SNX1 retained its punctate pattern in all samples, although we observed significant perinuclear accumulation of the signal in SWIP KO cells. FAM21-EGFP localized to SNX1-positive endosomes in WT cells, and to a lesser extent in control SWIP KO cells. However, it displayed a diffuse cytosolic signal in SWIP KO siVPS35 cells, which highlights how FAM21 depends on the two anchors for its endosomal localization.

**Figure 4.**
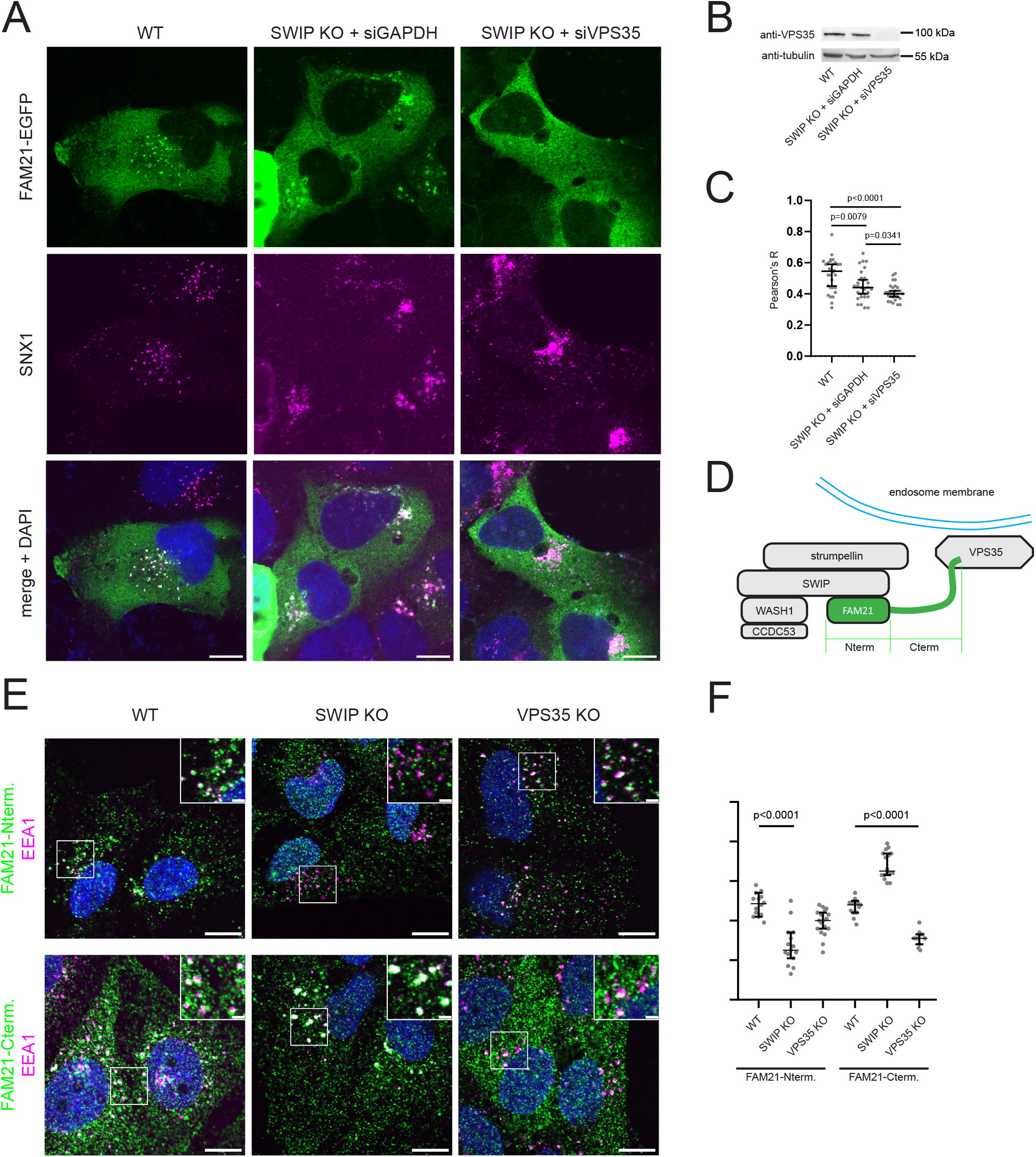
FAM21 binds to the endosomes via both SWIP and VPS35. **(A)** Cells were treated with the indicated siRNA for 24 h, transfected with FAM21-EGFP for further 24 h, fixed, and labeled with antibody targeting SNX1. **(B)** Western blot of wild-type U-2 OS cells and SWIP KO line treated with siRNA targeting GAPDH or VPS35, stained with antibody targeting VPS35. **(C)** Quantification of co-localization between FAM21-EGFP and SNX1. Each dot represents one analyzed cell. Data from 3 independent experiments, >25 cells in total analyzed in each column (nuclear regions removed from analyzed ROIs). **(D)** Schematic representation of WASH complex and retromer at the endosomal membrane is shown for better orientation in the experiments of this figure. **(E)** N-terminal FAM21-myc or C-terminal FAM21-myc fragments were transduced into wild-type, SWIP KO, or VPS35 KO cell lines. Resulting stable cell lines were fixed and labeled with antibodies targeting myc tag (green) and EEA1 (magenta). **(F)** Quantification of co-localization between FAM21-myc fragments and EEA1 in stable cell lines. Each dot represents one analyzed cell. Data from 2 independent experiments, >10 cells in total analyzed in each column. Scale bars: 10 μm (main images), 2 μm (insets). Quantitative data in graphs tested for normality with Shapiro-Wilk test, significance calculated using the unpaired t-test. Lines in graphs indicate median ± 95% confidence interval.

To further confirm this, we analyzed the localization of WASH1 in SWIP KO siVPS35 cells. We already established that WASH1 levels do not decrease in our KO cell lines and that endosomal localization of WASH1 strictly depends on FAM21. WASH1 thus serves as an indicator of FAM21 localization. Indeed, WASH1 also delocalized from endosomes in cells depleted of both SWIP and VPS35 (Fig. S4C).

Since it is known that FAM21 binds into the WASH complex via its N-terminal globular head domain, while the interaction with VPS35 is mediated by its C-terminal unstructured tail (Gomez and Billadeau, 2009), we investigated the localization of these FAM21 fragments (Fig. 4D) in WT, SWIP KO, and VPS35 KO cells. We expressed both domains as myc-tagged constructs using retroviral transduction to achieve low expression levels comparable to endogenous levels (Fig. 4E–F). Both fragments are localized to endosomes in WT cells. Indeed, the N-terminal globular domain localizes to endosomes in VPS35 KO cells, but not in SWIP KO cells. The opposite trend was apparent with the C-terminal tail, which localized to endosomes in SWIP KO cells, but this localization was decreased in the VPS35 KO cell line. Residual membrane recruitment of the C-terminal domain seen in VPS35 KO cells may be explained by the known affinity of FAM21 to phospholipids (Jia et al., 2010) or to other membrane proteins. Increase in co-localization seen with the FAM21 C-terminal domain in SWIP KO is a result of significant endosomal enlargement that leads to a higher overlap between the endosomal microdomains. Taken together, these data confirm that FAM21 is attached to the endosomal membrane via both SWIP and VPS35.

### SWIP binds to distinct phospholipid species in the endosomal membrane

We speculated that the endosomal anchoring of SWIP is mediated via binding to phospholipid species embedded in the membrane of the endosome. To address this, we attempted to purify individual WASH complex subunits from bacterial and insect expression systems. However, we encountered issues with protein solubility in the absence of their interaction partners, likely due to exposed hydrophobic surfaces where the subunits normally bind together. Nevertheless, we were able to purify a complete WASH complex from U-2 OS SWIP KO cells expressing EGFP-SWIP at near-endogenous levels. All five subunits were present in the purified sample (Fig. 5A), which we applied to a PIP strip membrane. The complete WASH complex binds strongly to PI, PI3P, PI4P, and PI5P and has medium affinity to PS and PI(3,5)P_2_ (Fig. 5B).

**Figure 5.**
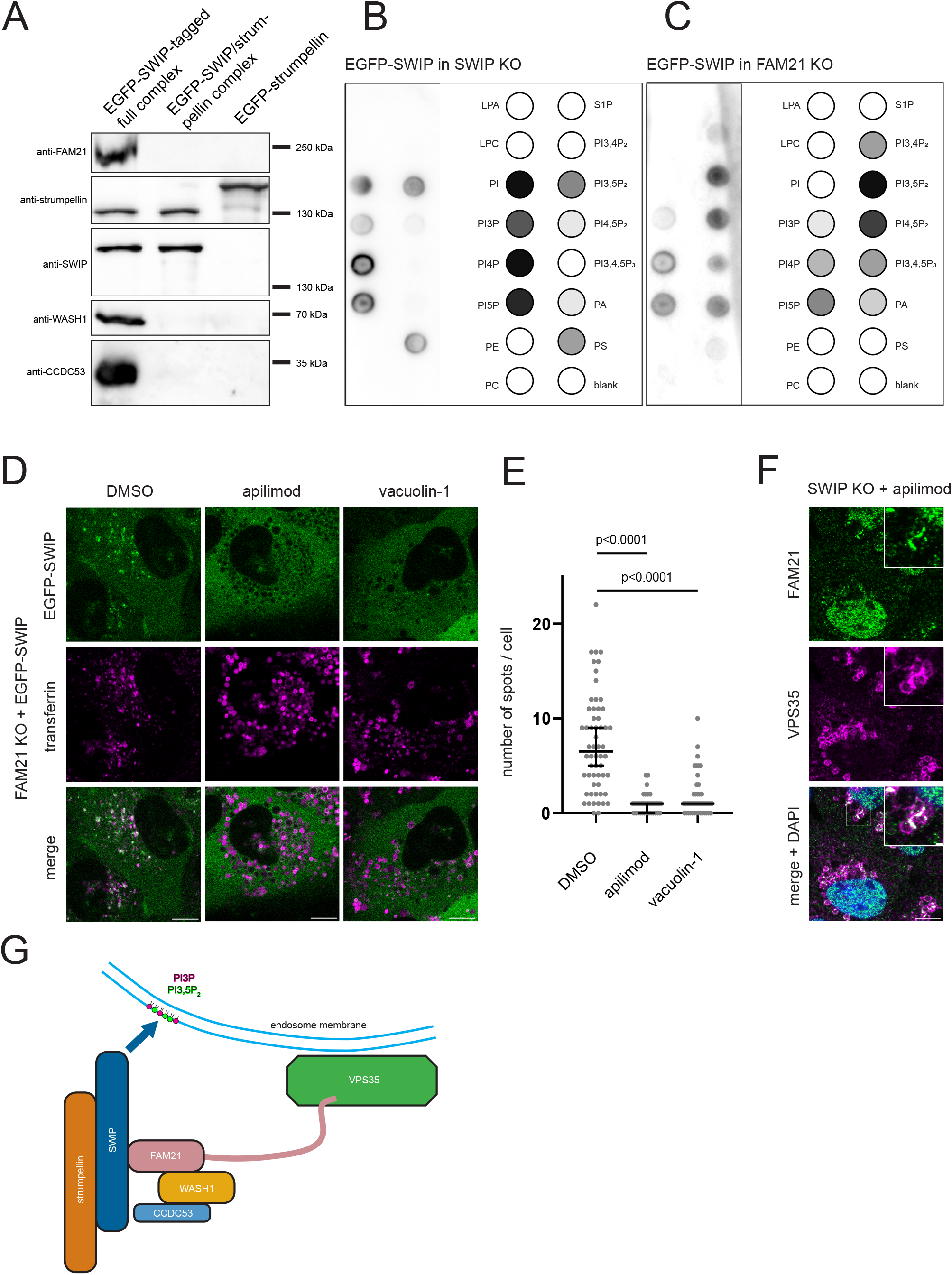
SWIP binds to distinct phospholipid species in the endosomal membrane. **(A)** SWIP KO cells expressing EGFP-SWIP, FAM21 KO cells expressing EGFP-SWIP and FAM21KO/SWIP KO (double KO) cells expressing EGFP-strumpellin were lysed and the WASH complex purified on GFP trap. Western blot incubated with indicated antibodies shows WASH subunits on beads (before elution) in an amount corresponding to the load on PIP strip. **(B)** PIP strip with WASH complex eluted from SWIP KO + EGFP-SWIP line, incubated with anti-SWIP antibody. One representative image shown. Schematic diagrams were prepared by averaging 3 independent experiments. **(C)** PIP strip with rudimentary WASH complex eluted from FAM21 KO + EGFP-SWIP line, incubated with anti-SWIP antibody. One representative image shown. Schematic diagrams were prepared by averaging 3 independent experiments. **(D)** FAM21 KO EGFP-SWIP cell line was treated with transferrin-Alexa 594 to label the endocytic compartment and then with DMSO, apilimod, or vacuolin-1 for 2 hours. Live confocal imaging without the fixation step. **(E)** Quantification of inhibitor treatments showing number of EGFP-SWIP spots per cell. Each dot represents one analyzed cell. Data from 3 independent experiments, >30 cells analyzed in each column. **(F)** SWIP KO cell line was treated as in (D), fixed, and stained with anti-FAM21 and anti-VPS35 antibodies. **(G)** Updated schematic model of WASH complex membrane attachment, depicting both retromer- and SWIP-mediated interaction with endosomal surface. Scale bars: 10 μm (main images), 2 μm (insets). Quantitative data in graph tested for normality with Shapiro-Wilk test, significance calculated using the Mann-Whitney test because of non-Gaussian distribution in the samples. Lines in graphs indicate median ± 95% confidence interval.

We wondered which of the identified interactions could be attributed to FAM21 and which to SWIP, as FAM21 was reported to bind several phospholipid species (Jia et al., 2010). Attempts to purify SWIP from cells lacking all other WASH complex subunits were unsuccessful as the cell line was not viable. We therefore resorted to KO cell lines lacking either FAM21 or both FAM21 and SWIP (double KO). We then purified the partial complex via one of the residual subunits tagged with EGFP. Immunoprecipitation of EGFP-SWIP from FAM21 KO yielded a complex consisting of SWIP and strumpellin, while immunoprecipitation of EGFP-strumpellin from double KO cells yielded only strumpellin (Fig. 5A). We reasoned that all phospholipid interactions detected with the SWIP-strumpellin complex, but not with strumpellin, can be attributed to SWIP. The SWIP-strumpellin complex showed strongest binding to PI bisphosphates, namely PI(3,5)P_2_ and PI(4,5)P_2_, but also had weaker affinity to PI(3,4)P_2_, PI(3,4,5)P_3_, PI3P, PI4P, PI5P, and phosphatidic acid (PA) (Fig. 5C). Contrary to that, we could not demonstrate any phospholipid binding of strumpellin from double KO cells, suggesting that it does not have detectable phospholipid-binding activity under conditions tested. Taken together, the observed phospholipid interactions are likely intrinsic to the SWIP subunit.

Phosphoinositide binding assayed with a PIP strip should be further confirmed with biological *in situ* experiments to show that the observed association is relevant in the specific intracellular context. In this respect, we considered PI3P and PI(3,5)P_2_ as the most obvious candidates since they are known as hallmarks of the endosomal membrane (with prominent enrichment on early and late endosomes, respectively). We used the FAM21 KO cell line stably expressing EGFP-SWIP, which cannot bind to membranes via FAM21 or VPS35, as a tool to assay SWIP-dependent WASH complex membrane binding. Contribution of PI3P and PI(3,5)P_2_ can be elucidated from the effects of PI3 kinase and PIKfyve inhibitors: if decreased recruitment of SWIP was observed upon treatment with PI3 kinase inhibitors alone, it would suggest a role of PI3P in binding. If SWIP bound to PI(3,5)P_2_, it should react sensitively to both PI3 kinase and PIKfyve inhibitors, because both reduce cellular levels of PI(3,5)P_2_ (Zhang et al., 2007). To test these hypotheses, FAM21 KO EGFP-SWIP cell line was pre-loaded with fluorescently labeled transferrin to visualize the endocytic compartment and then treated with inhibitors of PI3 class III kinase (SAR405) and PIKfyve (vacuolin-1, apilimod). In all cases, treatment leads to almost complete dissociation of SWIP from endosomal membranes (Fig. 5D and S5A), suggesting a prominent role of PI(3,5)P_2_ in SWIP-dependent WASH complex recruitment. Enormous change in vesicular morphology elicited by the inhibitors prevented a classical colocalization analysis. Therefore, number of SWIP-positive vesicles per cell was used for quantification (Fig. 5E and S5B). Sensitivity to PI(3,5)P_2_ is specific to SWIP, as FAM21 still localizes to vesicles in a SWIP KO cell line treated with apilimod (Fig. 5F).

We therefore propose an updated model of the WASH complex recruitment to the endosomal membrane (Fig. 5G). The complex features two main anchors – FAM21 and SWIP – attaching it to the vesicular surface. Absence of either of them leads to partial detachment of the complex from the membrane. The SWIP anchor is mediated via phosphoinositides in the endosomal membrane, exemplified by PI(3,5)P_2_, the removal of which leads to dissociation of SWIP from endosomes.

## Discussion

In recent years, accumulating evidence has shown that the WASH complex binds to membranes in both a retromer-dependent and retromer-independent fashion. Understanding of the process is confounded by a plethora of known interaction partners and the pentameric nature of the WASH complex, with each subunit potentially implicated in the membrane recruitment. Previous studies characterized mammalian gene knockouts of WASH1 (Gomez et al., 2012; Piotrowski et al., 2013), strumpellin (Tyrrell et al., 2016), and FAM21 subunits (McNally et al., 2017) and uncovered their various functional aspects. However, none of the reported functions explain WASH complex recruitment to the endosomal membrane.

This is the first comprehensive probe into WASH complex membrane binding, using a set of four subunit knockouts of the WASH complex itself as well as a VPS35 knockout for the retromer complex. We show that retromer depletion on its own does not prevent membrane binding of the WASH complex or hinder its endosomal role in F-actin nucleation. This has led us to hypothesize that the WASH complex uses other intrinsic membrane anchors. Conflicting data on the retromer essentiality for the endosomal recruitment of the WASH complex (McGough et al., 2014; Zavodszky et al., 2014) may be explained by use of different model systems, conditions, and cell types, each with a different demand for the retromer-based membrane anchor.

We speculate that a fraction of the WASH complex in cells may work without the retromer, as neither the attachment of the complex or the resulting F-actin polymerization are strictly dependent on the retromer. Furthermore, our results may explain how the retriever associates with endosomal membranes. This retromer-related complex requires the WASH complex for its membrane recruitment, while depletion of the retromer does not lead to a dissociation of the retriever (McNally et al., 2017). We speculate that this may be due to intrinsic membrane binding of the WASH complex itself.

Our work adds further background for the role of the Parkinson’s disease-linked VPS35 mutation D620N, which has previously been shown to reduce association between the retromer and the WASH complex (Cui et al., 2021; McGough et al., 2014). Given that the WASH complex associates with endosomes in the absence of VPS35, the mutation likely tunes the coordination between two membrane complexes – the retromer and the WASH complex – which are already membrane-associated independently of each other. The substantial autonomy of the two complexes is further documented by our observation that both N-terminal and C-terminal domains of FAM21 associate with endosomes. While previous studies indicated that the N-terminal globular domain of FAM21 binds to the rest of the WASH complex, a seemingly cytosolic distribution was observed whenever the C-terminal tail was removed from FAM21 (Jia et al., 2012). We show that the endosomal localization of the N-terminal domain becomes apparent when expressed at sufficiently low levels. Moreover, the N-terminal domain of FAM21 retains its endosomal localization in cells depleted of VPS35, while the C-terminal domain requires VPS35.

Our experiments show that SWIP is sufficient for endosomal binding of the complex even in the absence of FAM21. SWIP is an understudied component of the WASH complex with mostly helical arrangement and few structural domains that could provide insights into its function. Intriguingly, the P1019R substitution in SWIP is implicated in one form of autosomal recessive intellectual disability (Ropers et al., 2011), and the mutation leads to decreased affinity of SWIP to other WASH complex subunits (Courtland et al., 2021). Understanding the membrane-binding role of SWIP in the pathogenesis of this form of intellectual disability merits further research. Additionally, SWIP is known to be sequence-related to the WAVE complex subunit CYFIP1/2 (also known as Sra1), which has also been linked to congenital mental retardation (Abekhoukh and Bardoni, 2014; Jia et al., 2010). Similar to SWIP, CYFIP serves as one of the two main membrane anchors of the WAVE complex (Kurisu and Takenawa, 2009). However, this is not mediated by phosphoinositides but instead by the direct binding of CYFIP to small GTPase Rac (Schaks et al., 2018). A report of CYFIP1/2 directly binding to acidic phospholipids (Tsujita et al., 2010) has since not been confirmed.

The essential role of SWIP within the WASH complex was previously documented in the slime mold *Dictyostelium*, where knockout of SWIP, but not FAM21 or any other WASH complex subunit causes complete delocalization of GFP-WASH from lysosomal membrane (Park et al., 2013). These results reinforce the view that the WASH complex of primitive eukaryotes has a very close structural and functional resemblance to that of humans, despite limited sequence similarity (Seaman et al., 2013). On the other hand, we do not observe hyper-activated vesicular F-actin assembly in FAM21 KO cells, as is the case in *Dictyostelium* (Park et al., 2013), and our results suggest that three out of four major WASH complex subunits (WASH1, FAM21 and SWIP) are essential for WASH1 function in the U-2 OS human cell line. As an exception, the strumpellin KO cell line features weak F-actin spots on endosomes in some cells. This points to a partial redundancy of strumpellin for the function of the whole complex, which is in concordance with a recent work in the MDA-MB-231 human cell line (Tyrrell et al., 2016).

SWIP binding to the membrane likely occurs via its affinity to one or several phospholipid species. Among those identified to be bound by SWIP, PI(3,5)P_2_ and PI3P were our primary candidates from the PIP strip overlay assay. Our results show that SWIP detaches from endosomes in response to inhibitors of both PI3 kinase and PIKfyve, which points to PI(3,5)P_2_ as the most likely binding partner, because both classes of inhibitors are known to reduce PI(3,5)P_2_ levels. Indeed, inhibition of PI3P production was recently shown to decrease FAM21 localization to VPS35-positive vesicles (Giridharan et al., 2022). In the same work, decrease in PI(3,5)P_2_ did not elicit WASH complex detachment. However, our experiments were conducted on a FAM21 KO cell line and thus shed light on the phosphoinositide-binding capability of SWIP rather than the whole WASH complex. We show that the FAM21 anchor is not sensitive to apilimod treatment, in contrast with SWIP. We hypothesize that SWIP could assist FAM21 in attaching the WASH complex to endosomes in the increasingly PI(3,5)P_2_-rich late endosomal compartment, but this requires further research.

The membrane affinity of SWIP could be a property of the protein itself. However, SWIP has no lipid-binding domains. Alternatively, its binding could also be mediated indirectly by a SWIP-interacting protein outside the WASH complex. To our knowledge, a viable candidate for such a protein has not been described yet. Additionally, we cannot rule out other phosphoinositides being at play in the membrane attachment of SWIP. Indeed, SWIP has some affinity towards PI4P, PI5P, and PI(3,4)P_2_ in the PIP strip overlay assay, and all these phosphoinositides were implicated in the regulation of endosomal traffic (Boal et al., 2015; Dong et al., 2016; Feng and Yu, 2021).

Attachment of the WASH complex to endosomes is essential for its role in recruitment and activation of a local pool of Arp2/3. We have elucidated a novel contact between the WASH complex subunit SWIP and the endosomal surface which contributes to this important process besides the canonical retromer link. This SWIP-mediated anchor may increase the specificity or strength of the sub-endosomal localization of the WASH complex. Further research will hopefully shed more light on the structural mechanism of SWIP membrane recruitment and the way in which this function of SWIP is reflected in genetic diseases associated with the protein and its interactome.

## Materials and methods

### Plasmids and siRNA

For the expression of N-terminal and C-terminal fragments of FAM21, 1-354aa and 355aa-end domains of FAM21 were PCR-amplified from human U-2 OS cDNA and inserted into pcDNA4TOMycHis vector via HindIII and BamHI (N-terminal) or HindIII and ApaI (C-terminal). Retroviral pMX-Puro vectors pMX-FAM21^1-354aa^-myc and pMX-FAM21^355aa-end^-myc were prepared from pMX-puro vector by insertion of myc-tagged fragments at BclI and XhoI sites, which were PCR-amplified from respective pcDNA4TOMycHis vectors. Full FAM21 was PCR-amplified from human U-2 OS cDNA and inserted into pEGFP-N1 via XhoI and BamHI sites. Vector pMX-EGFP-SWIP was made by PCR amplification of human EGFP-SWIP from pEGFP-C1-SWIP and insertion into pMX-Puro via BamHI and NotI. For purification experiments, a PreScission site was added between EGFP and SWIP by site-directed PCR mutagenesis. SWIP sequence from EGFP-PreScission-SWIP was then replaced with strumpellin sequence (via Gibson assembly) to yield EGFP-PreScission-strumpellin. For pSpCas9(BB)-2A (PX459) vectors, oligos carrying the target CRISPR sequence were phosphorylated, annealed, and subsequently inserted into BpiI-digested PX459 backbone. All vectors were verified by sequencing before use.

Silencer Select Pre-designed siRNAs were ordered from Ambion. All primers and siRNAs used in this study are summarized in **Table S1** and **S2**.

### Cell culture and transfection

U-2 OS cells (ATCC) were cultured in Dulbecco’s modified Eagle’s medium (DMEM + GlutaMAX, Gibco) supplemented with 10% fetal bovine serum and 100 U/mL penicillin-streptomycin solution (Biowest). Cells were authenticated at the start of the project and regularly inspected for mycoplasma contamination. For long-term storage at −80 °C, cells were frozen in DMEM with 10% serum and 10% dimethyl sulfoxide (DMSO), or in DMEM with 20% KnockOut Serum Replacement (Thermo Fisher) and 10% DMSO for the FAM21 knockout cell line.

Transient DNA transfection of these cells was performed using X-tremeGENE HP DNA Transfection Reagent (Roche) according to manufacturer’s instructions. For transfection of siRNA, we used 1.5 μL Lipofectamine RNAiMAX (Invitrogen) and 10 pmol siRNA per 1 mL of cell culture medium and subsequently incubated with cells for 48 h. Cells were thensplit, seeded and the process was repeated for another 48 h to achieve complete silencing.

### Generation of KO and stable cell lines

U-2 OS knockout cell lines were generated by CRISPR/Cas9 using vector pSpCas9(BB)-2A (PX459) as described (Ran et al., 2013) with sequence-specific oligos (**Table S3**) selected from Gecko v2 Knockout Library (Sanjana et al., 2014). U-2 OS cells were transiently transfected with a single pSpCas9 vector (*strumpellin*) or a 1:1 mixture of two pSpCas9 vectors targeting different parts of the gene of interest (*VPS35, WASH1, FAM21*). Cells were selected with 2.5 μg/mL puromycin for 3 days and single cell clones were generated. Cell line “Double KO” was generated from FAM21 KO by knocking out *SWIP* using identical procedures. For the SWIP KO cell line, sgDNA was first PCR-amplified from pDD162 vector (Addgene 47549) using primers targeting the *SWIP* sequence. The PCR products were transcribed with T7 HiScribe kit (New England Biolabs) and transfected together with TrueCas9 protein v2 (Thermo Fisher) and Lipofectamine CRISPRmax reagent (Thermo Fisher). All single-gene knockout cell lines were verified using SDS PAGE/Western blot and sequencing of the affected regions of the genes. Cell lines used in this work are summarized in **Table S4**.

For retroviral stable cell lines, pMX-Puro vectors were transfected into Platinum-A packaging line (Cell Biolabs) grown in standard DMEM medium with 10% serum. Medium with the viral titer was harvested after 3 days, centrifuged at 500g for 10 min to remove detached Platinum-A cells, and the resulting supernatant was applied to target cell line. Medium was exchanged after 24 h and cells were selected with 2.5 μg/mL puromycin for 3 days to yield polyclonal stable cell lines.

### Inhibitor treatments

For treatment with inhibitors, cells pre-labeled with transferrin-Alexa Fluor 594 were treated for 2 hours with 10 μM SAR405 (Merck, cat. no. 5330630001; dissolved in DMSO), 1 μM apilimod (Selleckchem, cat. no. S6414; dissolved in DMSO), 10 μM vacuolin-1 (Merck, cat. no. 673000; dissolved in DMSO), or a corresponding volume of DMSO as a control and then directly processed for live microscopy.

### Antibodies

The following primary antibodies were used for Western blotting (WB) and immunofluorescence (IF): anti-cMyc (Exbio, cat. no. 11-433-C100, 1:1000 for IF), anti-alpha-tubulin (Sigma-Aldrich, cat. no. T9026, 1:10000 for WB), anti-VPS35 (Abcam, cat. no. ab10099, 1:200 for IF, 1:1000 for WB), anti-WASH1 (Atlas Antibodies, cat. no. HPA002689, 1:200 for IF, 1:1000 for WB), anti-strumpellin (Abcam, cat. no. ab101222, 1:400 for IF, 1:1000 for WB), anti-SWIP (Bethyl Laboratories, cat. no. A304-919A-M, 1:1000 for IF and WB), anti-FAM21 (Santa Cruz, cat. no. sc-137995, 1:200 for IF, 1:1000 for WB), anti-CCDC53 (Atlas Antibodies, cat. no. HPA038339, 1:200 for IF), anti-CCDC53 (Millipore, cat. no. ABT69, 1:1000 for WB), anti-EEA1 (Cell Signaling, cat. no. 3288, 1:800 for IF), anti-SNX1 (Santa Cruz, cat. no. sc-136247, 1:200 for IF) and anti-GLUT1 (Abcam, cat. no. ab115730, 1:400 for IF).

Secondary antibodies used for Western blot were Horseradish peroxidase goat antimouse (Jackson ImmunoResearch, cat. no. 115-035-146, 1:10000), Horseradish Peroxidase goat anti-rabbit (Jackson ImmunoResearch, cat. no. 111-035-045, 1:10000) and Horseradish peroxidase bovine anti-goat (Jackson ImmunoResearch, cat. no. 805-035-180, 1:5000). For immunofluorescence, goat anti-mouse IgG (H+L) secondary antibody-Alexa Fluor 488 (Thermo Fisher, cat. no. A11001) was used together with donkey anti-rabbit IgG (H+L) secondary antibody-Cy5 (Jackson ImmunoResearch, cat. no. 711-175-152, 1:500). Alternatively, donkey anti-rabbit IgG (H+L) secondary antibody-Alexa 488 (Jackson ImmunoResearch, cat. no. 711-545-152, 1:500) was used together with transferrin-Alexa Fluor 594 or with donkey anti-goat IgG (H+L) secondary antibody-Alexa 594 (Thermo Fisher, cat. no. A-11058, 1:500).

### Immunofluorescence and microscopy

Cells were grown on glass coverslips and treated as indicated. For transferrin staining, cells were first serum starved for 1.5 h in DMEM, treated with 5 μg/mL transferrin-Alexa Fluor 594 (Invitrogen) for 15 min, and then further processed. Cells were fixed for 10 minutes in 3% paraformaldehyde solution (Sigma-Aldrich) in microtubule-stabilizing buffer (20 mM 2-N-Morpholino-ethanesulfonic acid, 2 mM EGTA, 2 mM magnesium chloride, 4% w/v PEG 6000, 137 mM sodium chloride, 5 mM potassium chloride, 1.1 mM dibasic sodium phosphate, 0.4 mM monopotassium phosphate, 5.5 mM glucose, 4 mM sodium bicarbonate).

The coverslips were then washed 3 × 5 minutes in PBS, and cells were permeabilized in 0.1% Triton X-100 (Sigma-Aldrich) in PBS for 5 minutes and washed again. Coverslips were then incubated with primary antibodies (diluted in 3% bovine serum albumin in PBS) for 45 minutes and washed again. Finally, cells were incubated in secondary antibodies (diluted in 3% bovine serum albumin in PBS) for 45 minutes and washed again. For double immunofluorescence, primary antibodies were used sequentially while the secondary antibody staining was simultaneous. For actin staining, Alexa Fluor 488 phalloidin (Molecular Probes, 1 μL per 22×22 mm coverslip) was added to the secondary antibody step. All coverslips were then mounted onto Mowiol supplemented with 0.1 μg/mL DAPI (Sigma-Aldrich).

Cells were observed on confocal microscope (Leica TCS SP8) with HC PL APO 63×/1.40 oil immersion objective and HyD and PMT detectors. HyD was preferred for detection when possible. Images were acquired from LASX software at 16bit depth and prepared for publication in ImageJ Fiji. Images in figures 4E, 5D, 5F, and S5A were acquired on a Zeiss LSM 880 confocal microscope with PL APO 63×/1.40 oil immersion objective and PMT detector, exported from Zeiss Zen, and further processed as described above.

Colocalization calculations (Pearson’s R without threshold) were executed in Coloc 2 plugin for ImageJ Fiji. Where indicated, nuclear regions were not included in the analyzed ROI because of potential variations in levels of unspecific nuclear signal between experiments.

### SDS PAGE and Western blotting

Cell pellets for SDS polyacrylamide gel electrophoresis were dissolved in sample buffer (62.5 mM Tris pH 6.8, 2% SDS, 10% glycerol, 5% 2-mercaptoethanol, traces of bromophenol blue), heated at 95 °C for 5 minutes, and centrifuged at 15,000 g for 5 minutes to remove insoluble debris. For electrophoretic separation, 8% polyacrylamide gel was used with 5% stacking layer. Gels were then transferred onto nitrocellulose membrane (pore size 0.2 μm, GE Healthcare) in wet conditions using methanol-based blotting buffer (25 mM Tris, 192 mM glycine, 20% methanol). Membranes were blocked in 3% skimmed milk in TBST for 1 hour and incubated overnight with primary antibody in 3% bovine serum albumin in TBST. Next morning, membranes were thoroughly washed and incubated with HRP-conjugated secondary antibody in 3% skimmed milk in TBST. Signal was developed with Pierce ECL substrate (Thermo Fisher) using LAS 4000.

### Protein purification and PIP strip overlay assay

SWIP and strumpellin were purified from corresponding retroviral U-2 OS cell lines stably expressing EGFP-SWIP or EGFP-strumpellin with PreScission protease sites after EGFP tag. Approximately 6×10^7^ cells were harvested by trypsin, lysed in 5 mL lysis buffer (50 mM HEPES pH 7.4, 100 mM KCl, 1 mM EGTA, 1 mM MgCl_2_, 2% NP-40), homogenized by 10 passes through a 26-gauge syringe, briefly sonicated, and centrifuged at 16,000 g for 20 minutes to remove cell debris. Cleared lysates were diluted with 5 mL dilution buffer (lysis buffer without NP-40) and incubated with 10 μL bead slurry (GFP-Trap Magnetic Particles M-270, Chromotek) for 2 h. Beads were then washed 5 times with dilution buffer and proteins were cleaved with 2 μg PreScission protease (MPI-CBG) in cleavage buffer (50 mM Tris-HCl, pH 7.0, 150 mM NaCl, 1 mM EDTA, 1 mM DTT) at 4 °C for 16 h. PIP Strips Membranes (Thermo Fisher) were blocked in 3% milk in TBST for 1 h, incubated with either purified SWIP or strumpellin diluted in 3% BSA in TBST (pH 7.4) overnight at 4 °C, washed, and then processed as described for Western blot membranes.

## Acknowledgements

The authors declare no competing financial interests. This work was funded by Charles University Grant Agency (GA UK, project number 1311920), Progress Q43, and SVV260559. Microscopy was performed in the Laboratory of Confocal and Fluorescence Microscopy co-financed by the European Regional Development Fund and the state budget of the Czech Republic, projects no. CZ.1.05/4.1.00/16.0347 and CZ.2.16/3.1.00/21515, and supported by the Czech-BioImaging large RI project LM2018129. Computational resources were supplied by the project “e-Infrastruktura CZ” (e-INFRA LM2018140) provided within the program Projects of Large Research, Development, and Innovations Infrastructures. We also thank Vladimír Kořínek for assistance with CRISPR and retroviral cell lines, Daniel Rozbeský and Tomáš Zdobinský for assistance with protein purification, and Jan Černý for vacuolin-1.

## Author contributions

**Vojtěch Dostál**: Conceptualization, Methodology, Formal analysis, Investigation, Visualization, Writing – Original Draft. **Tereza Humhalová**: Conceptualization, Methodology, Formal analysis, Investigation, Visualization, Writing – Original Draft, Funding acquisition. **Pavla Beránková**: Investigation. **Ondřej Pácalt**: Investigation. **Lenka Libusová**: Conceptualization, Methodology, Writing – Review & Editing, Supervision, Project administration, Funding acquisition.

