## Supplementary Data for "SWIP mediates retromer-independent membrane recruitment of the WASH complex"

### Supplementary figure 1

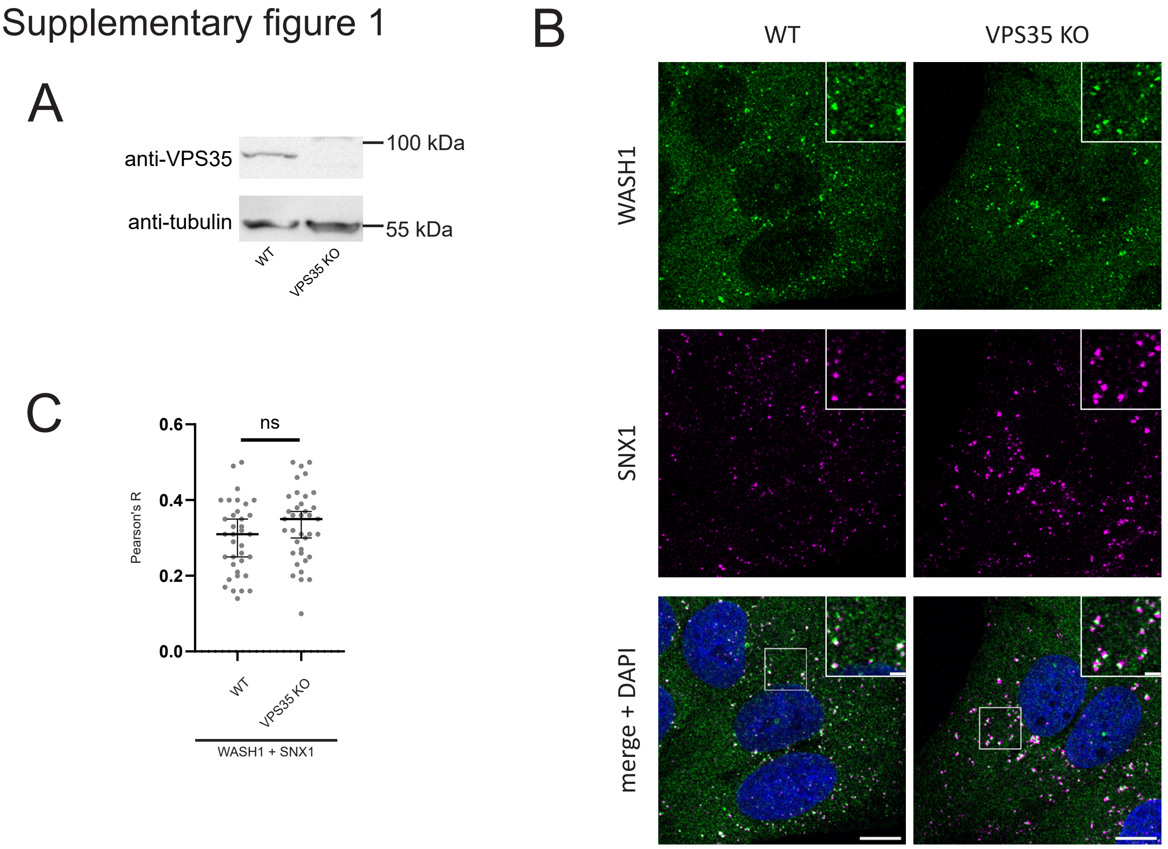


**(A)** Western blot confirming the absence of VPS35 in the VPS35 KO cell line. **(B)** Wild-type U-2 OS cells and VPS35 knockout U-2 OS cells were fixed and labeled with antibodies targeting SNX1 and WASH complex subunit WASH1. **(C)** Quantification of colocalization between SNX1 and WASH1 signal in WT and VPS35 KO cells. Each dot represents one analyzed cell. Data from 3 independent experiments, >20 cells in total analyzed in each column (nuclear regions removed from analyzed ROIs). Datasets tested for normality with Shapiro-Wilk test, significance calculated using the unpaired t-test. Lines in graph indicate median ± 95% confidence interval.

### Supplementary figure 2

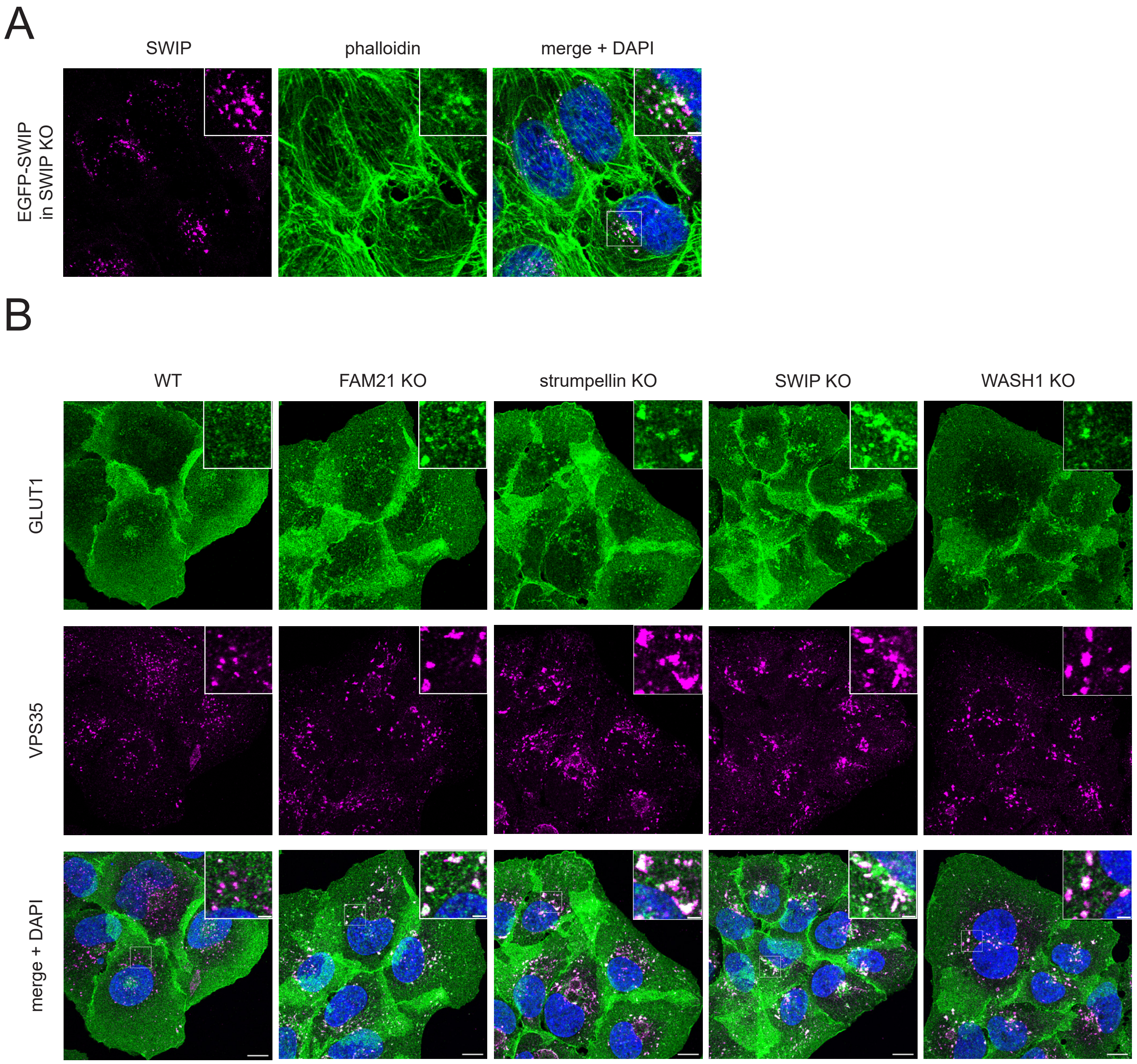


**(A)** Stable cell line with EGFP-SWIP expressed in SWIP KO, showing rescue of WASH function. Cells were fixed and stained with phalloidin to reveal endosomal actin patches.
**(B)** Wild-type U-2 OS cells and indicated knockout cell lines were fixed and labeled with antibodies targeting VPS35 and GLUT1. GLUT1 staining shows that the protein is sequestered to endosomes in FAM21 KO, strumpellin KO, SWIP KO and WASH1 KO cell lines.

Scale bars: 10 µm (main images), 2 µm (insets).

### Supplementary figure 3

A

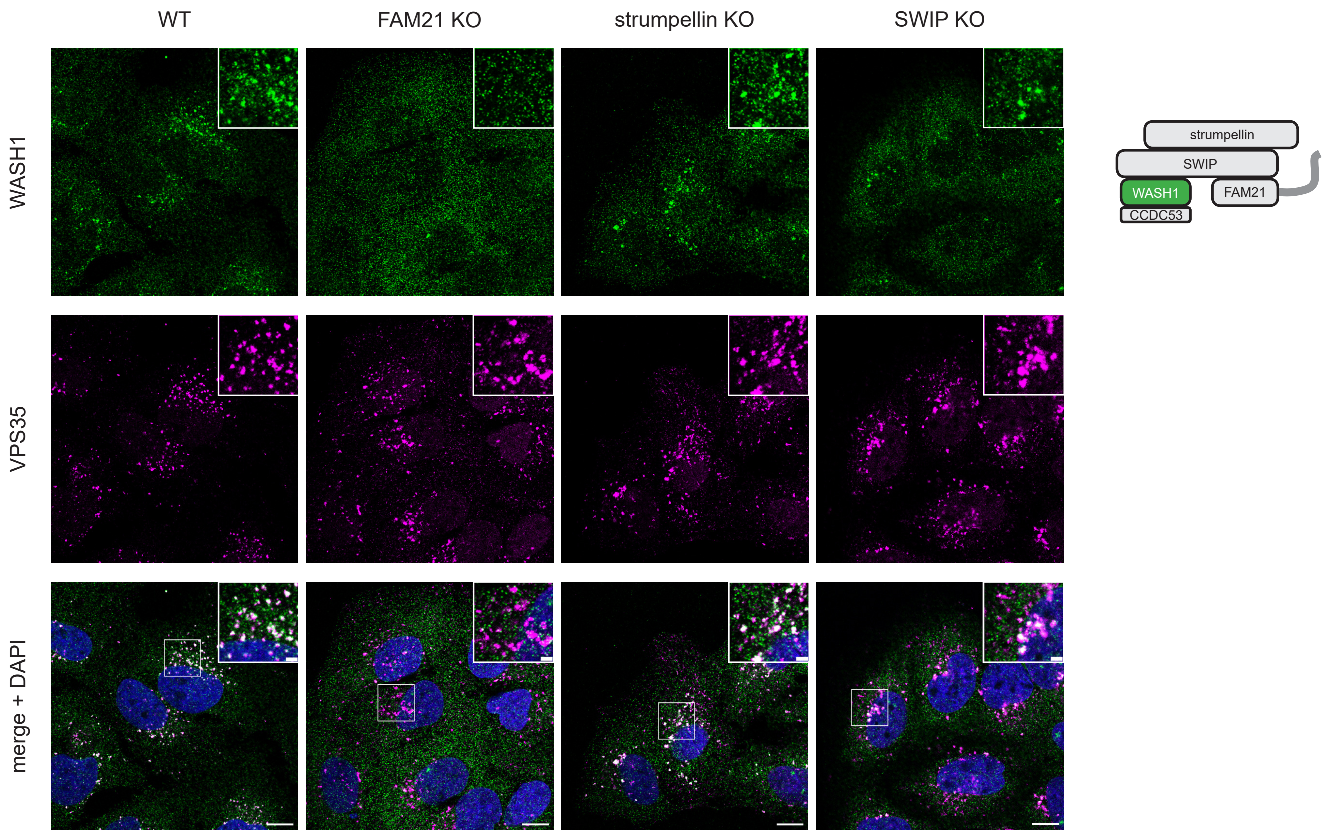


B

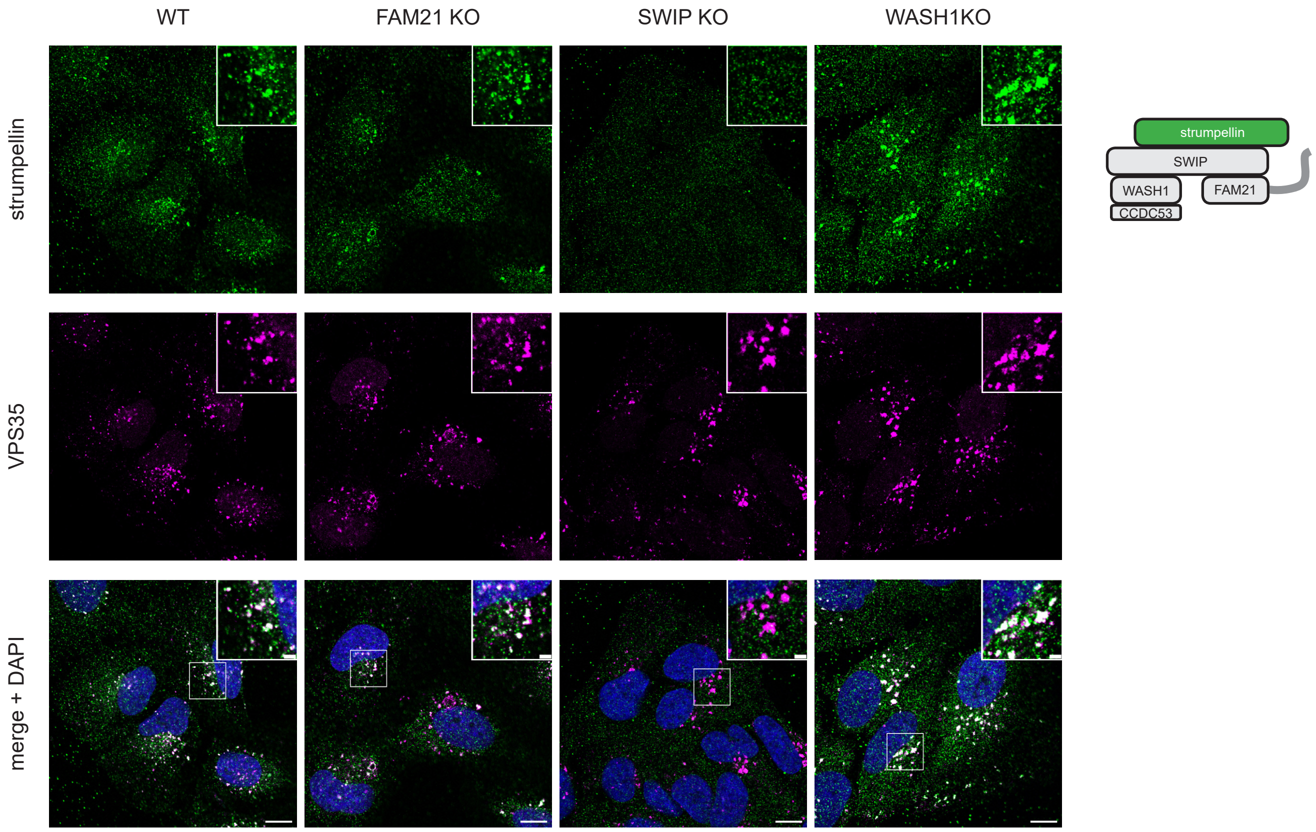

C

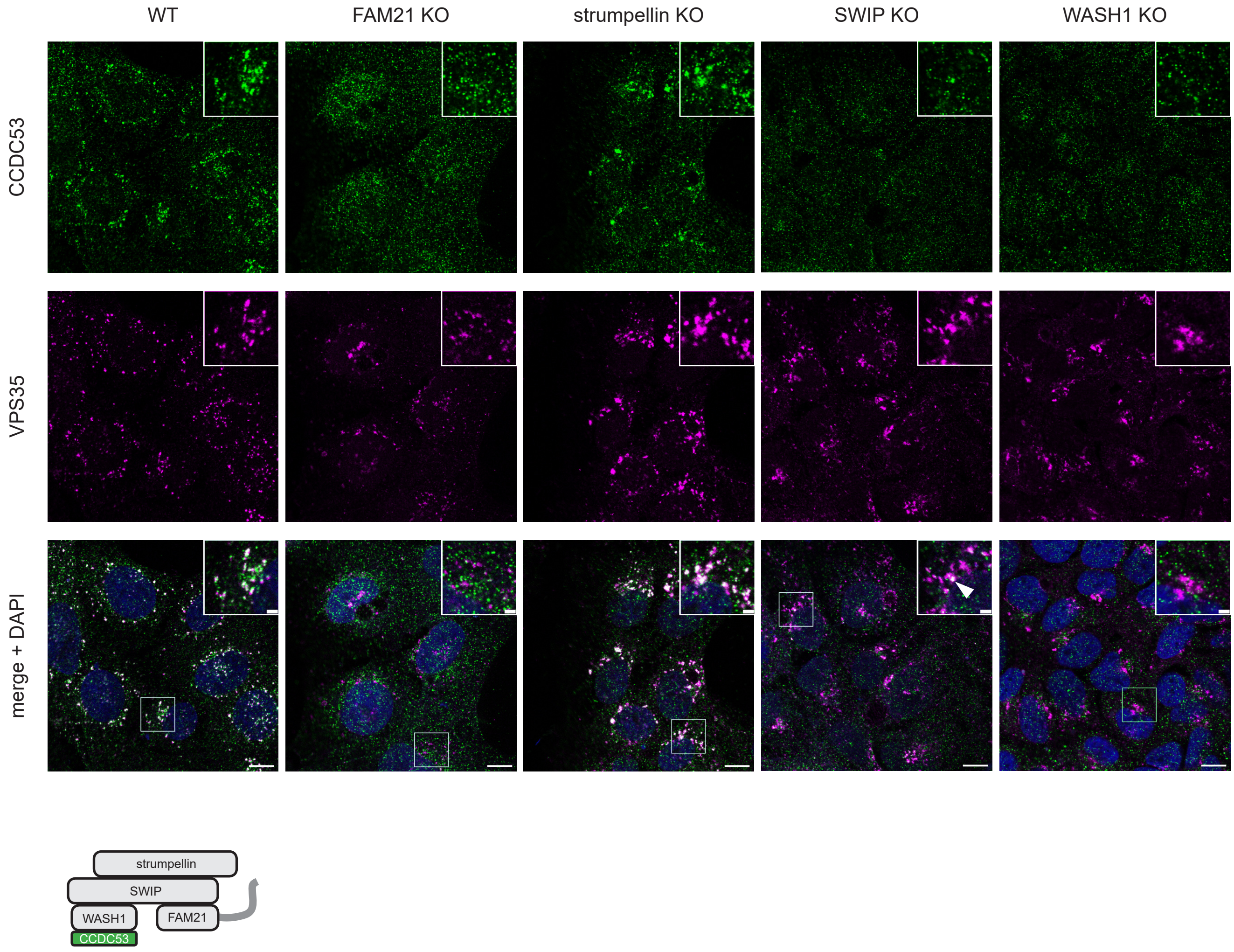


Wild-type U-2 OS cells and indicated knockout cell lines were fixed and labeled with antibodies targeting VPS35 and following proteins: WASH1 **(A)**, strumpellin **(B)** or CCDC53 **(C)**. Please note the delocalization of strumpellin in SWIP KO cell line, and delocalization of WASH1 and CCDC53 in FAM21 KO cell line. White arrow denotes CCDC53 signal on an endosome in the SWIP KO cell line.

Scale bars: 10 µm (main images), 2 µm (insets).

Supplementary figure 4

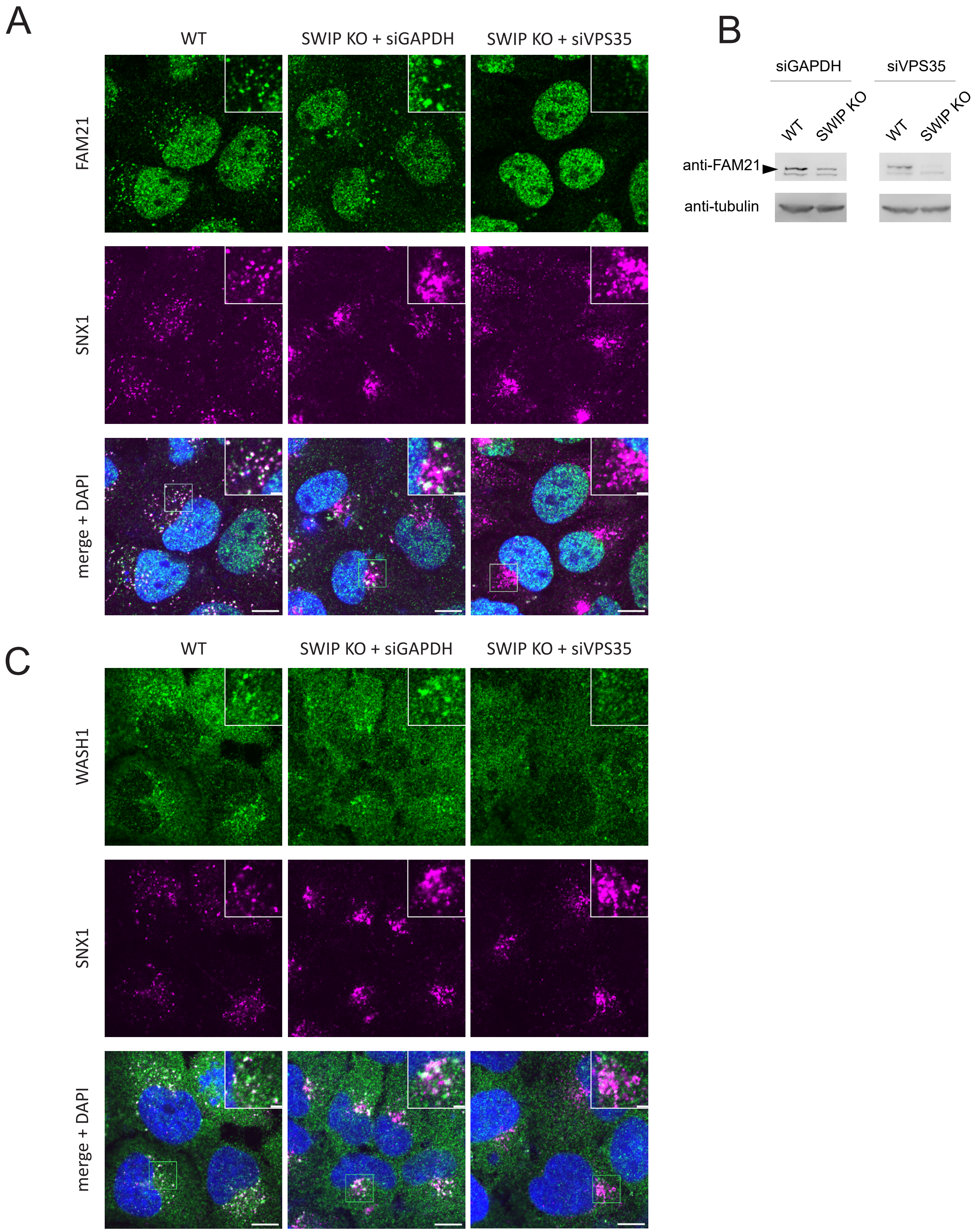


**(A)** Wild-type U-2 OS cells and SWIP KO cells treated either with control GAPDH siRNA (siGAPDH) or with VPS35 siRNA (siVPS35) were fixed and labeled with antibodies targeting FAM21 and SNX1.
**(B)** Western blot showing the decrease of the FAM21 protein levels in SWIP KO cells treated with siRNA against VPS35. Tubulin served as a loading control.
**(C)** Wild-type U-2 OS cells and SWIP KO cells treated either with control GAPDH siRNA (siGAPDH) or with VPS35 siRNA (siVPS35) were fixed and labeled with antibodies against WASH1 and SNX1.

Scale bars: 10 µm (main images), 2 µm (insets).

Supplementary figure 5

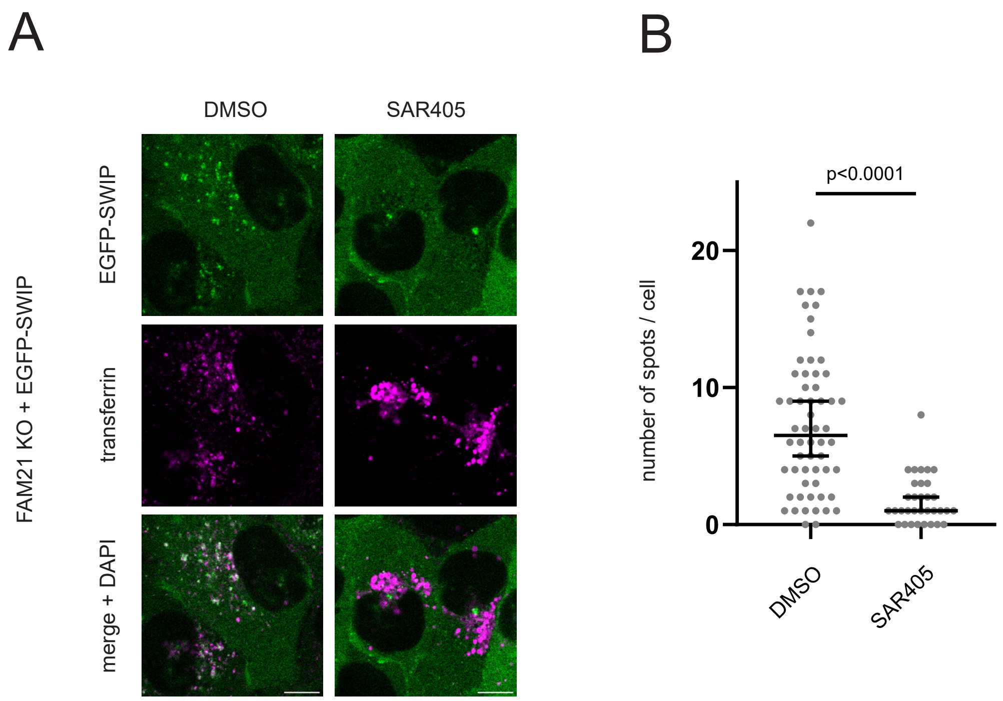


**(A)** FAM21KO EGFP-SWIP cell line was treated with transferrin-Alexa 594 and then with DMSO or SAR405. **(B)** Quantification of inhibitor treatments showing number of EGFP-SWIP spots per cell. DMSO data set identical to that in Fig. 5E because all inhibitor treatments were run in parallel. Data from 3 independent experiments, >30 cells analyzed in each column.

Scale bars: 10 µm. Quantitative data in the graph tested for normality with Shapiro-Wilk test, significance calculated using the Mann-Whitney test because of non-Gaussian distribution in the samples. Lines in graphs indicate median ± 95% confidence interval.
